# Topological insights into the neural basis of flexible behavior

**DOI:** 10.1101/2021.09.24.461717

**Authors:** Tevin C. Rouse, Amy M. Ni, Chengcheng Huang, Marlene R. Cohen

**Author notes:** TCR and MRC designed the project; TCR developed the analysis techniques and analyzed the data and simulations; AMN collected the electrophysiological data; CH performed the model simulations; MRC supervised the project; TCR, AMN, CH, and MRC contributed to writing the manuscript.

## Abstract

It is widely accepted that there is an inextricable link between neural computations, biological mechanisms, and behavior, but it is challenging to simultaneously relate all three. Here, we show that topological data analysis (TDA) provides an important bridge between these approaches to studying how brains mediate behavior. We demonstrate that cognitive processes change the topological description of the shared activity of populations of visual neurons. These topological changes constrain and distinguish between competing mechanistic models, are connected to subjects’ performance on a visual change detection task, and, via a link with network control theory, reveal a tradeoff between improving sensitivity to subtle visual stimulus changes and increasing the chance that the subject will stray off task. These connections provide a blueprint for using TDA to uncover the biological and computational mechanisms by which cognition affects behavior in health and disease.

**Significance Statement:** As the fields of systems, computational, and cognitive neuroscience strive to establish links between computations, biology, and behavior, there is an increasing need for an analysis framework to bridge levels of analysis. We demonstrate that topological data analysis (TDA) of the shared activity of populations of neurons provides that link. TDA allows us to distinguish between competing mechanistic models and to answer longstanding questions in cognitive neuroscience, such as why there is a tradeoff between visual sensitivity and staying on task. These results and analysis framework have applications to many systems within neuroscience and beyond.

---

**P**erhaps the most remarkable hallmark of the nervous system is its flexibility. Cognitive processes including visual attention have long been known to affect both behavior (e.g. performance on visual tasks) and virtually every measure of neural activity in visual cortex and beyond ((1),(2)). The diversity of changes associated with cognitive processes like attention makes it unsurprising that very simple, common measures of neural population activity provide limited accounts of how those neural changes affect behavior.

Arguably, the most promising simple link between sensory neurons and behavior is correlated variability (often quantified as noise or spike count correlations, or *r_SC_*, which measure correlations between trial-to-trial fluctuations in the responses of a pair of neurons to repeated presentations of the same stimulus; (3)). Correlated variability in visual cortex is related to the anatomical and functional relationships between neurons ((4); (3)). We demonstrated previously that the magnitude of correlated variability predicts performance on a difficult but simple visual task (Fig. 1D) across experimental sessions and on individual trials ((5)). This early success relating neural activity to simple behaviors means that correlated variability is a foundation on which to build efforts to explain more complex aspects of flexible behavior and the concomitant neural computations.

**Fig. 1.**
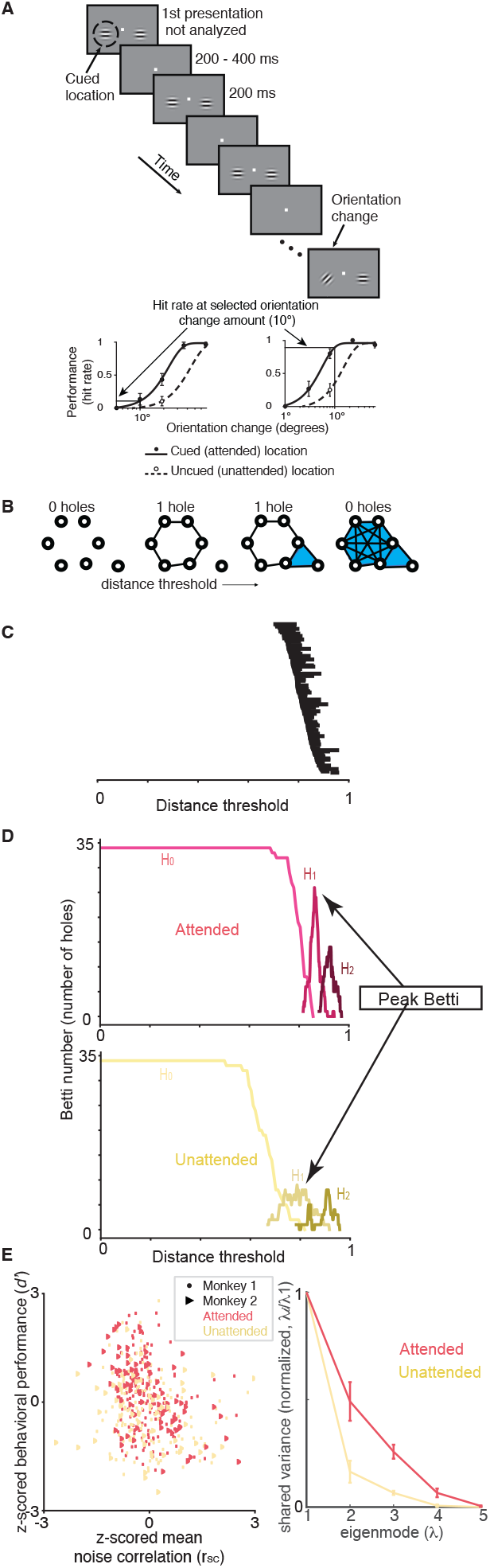
Experimental and topological methods. (A). Orientation change detection task with cued attention ((5)). The lower panels are psychometric curves (hit rate as a function of orientation change amount) for two example recording sessions to illustrate how we calculated performance at one selected orientation change amount on every recording session. (B). Illustration of topological data analysis methods. Each circle represents a neuron, and the distance between each pair is 1-their pairwise noise correlation (note that in real networks, more than two dimensions are typically required to represent all of the pairwise interactions). This analysis method iterates through distance thresholds (going from small to large from left to right). When the distance between two points is less than the threshold, they are considered connected. The shaded regions indicate groups of points that are fully interconnected, which indicates a higher order interaction between that subgroup of neurons. We summarize this structure by counting the number of holes at each threshold (which constructs the Betti curve in (D). For a more detailed description see Supplemental (Topological Data Analysis Example). (C). Topological description (persistence barcode) of an example recording session showing the distance thresholds (x-axis) at which each hole exists (holes are ordered by the threshold at which they appear). Many data sets are characterized by a small number of persistent topological features, which would show up as long horizontal lines in this plot. Instead, our neural data are characterized by a large number of holes that persist only for a small range of distance thresholds (many short horizontal lines in this plot). (D) Example Betti curves (plots of the number of holes as a function of distance threshold, which corresponds to the number of lines present at each threshold in (C)) for an example recording session. *H*_0_ (lightest color; *0th* dimension) curve keeps track of the number of connected components (‘holes’ equivalent to points) in the graph as the threshold is varied. *H*_1_ (middle color; 1*st* dimension) curve tracks the number of circular features with changes in the threshold. *H*_2_ (darkest color; 2*nd* dimension) curve tracks the number of spherical features with changes in the threshold. Our analyses focus on the peak of the Betti curve for the 1st and 2nd dimension (see supplement for other topological descriptions). (E) We focus our topological analyses of neural populations on correlated variability because there is a strong relationship between *r_SC_* and performance on our psychophysical task (quantified as sensitivity or *d*’). The plot shows *d*’ as a function of *r_SC_* (both values are z-scored for each animal and computed from responses to the stimulus before the orientation change). The colors represent the trials when attention was directed inside (‘attended’, red) or outside the receptive fields of the recorded V4 neurons (‘unattended’, yellow). The correlation between *d*’ and *r_SC_* was significant for each attention condition (attended: *r* = −0.11, *p* = 0.11; unattended *r* = −0.25, *p* = 4.58*e* – 5). (E) Factor analysis is a common linear method to assess the dimensionality of the correlated variability. The plot shows the shared variance (first five eigenvalues of the shared covariance matrix with private variance removed using Factor analysis) normalized by the shared variance in the first (dominant mode).

However, our efforts to relate correlated variability to a wider variety of sensory and cognitive phenomena and to constrain mechanistic models reveal a need for more sophisticated ways to characterize neuronal population activity. For example, although low correlations are associated with better performance in the case of attention and learning ((5); (6)), they are associated with worse performance when modulated by adaptation or contrast ((7)). Even in the case of cognitive processes like attention or task switching, good performance is associated with increases in correlation among particular subgroups of neurons ((8);(7)). And although mean correlated variability places much stronger constraints on cortical circuit models of cognition than measures of single neuron responses ((9); (10)), these models remain under-constrained.

These results highlight the need to use holistic methods to investigate the relationship between noise correlations and behavior. We focused on topological data analysis (TDA; (11); (12)), which is an emerging area in mathematics and data science that leverages groundbreaking advances in computational topology to summarize, visualize and discriminate complex data based on topological data summaries. These approaches, which have mostly been used in fields like astrophysics or large scale neural measurements (see, e.g., (13); (14); (15)), are able to identify features in the data that are qualitatively distinct from those highlighted using traditional analytic methods.

## 1. Results

### A. Topological signatures of correlated variability

We used TDA (specifically persistent homology; (16)) to quantify the higher-order structure in the pairwise interactions between simultaneously recorded neurons from area V4 of rhesus monkeys performing a difficult visual detection task with an attention cue (Figure 1A; different aspects of these data have been presented previously; (5)). We analyzed the structure of noise correlations in a population of neurons in the visual cortex by constructing a space in which the distance measure between a pair of neurons is 1 – *r_SC_*, where *r_SC_* is their noise correlation (Fig. 1B). In this space, highly correlated neurons are near each other, and anticorrelated neurons are far apart.

As is typical of the persistence homology approach, we iterate through a distance threshold (left to right in Fig. 1B) to understand the topological features of the correlation structure. For each threshold, we consider a pair of neurons to be functionally connected if their distance is less than the threshold. As the threshold increases, we thus include functional connections between pairs that are less correlated. For each distance threshold, we use established TDA methods to identify “holes” in the correlation structure, which correspond to the lack of connections between a subset of neurons and have implications for the organization and function of the network ((17), (18)).

We use TDA in a slightly different way than in most past work. The most common uses of TDA focus on persistent features (holes that persist through a large range of distance thresholds; (19); (20); (21); (22)). For example, a beautiful recent study used TDA to analyze the structure of signal represented by a population of neurons ((22)). Those authors focused on the persistent features of that data set, which reflect the quantities encoded by that population of neurons. In contrast, we analyzed noise, which is not thought to have any particular structure (much less one characterized by holes of different dimensionality). In our data sets, we simply did not observe persistent features (Figure 1C). Instead, we observed large numbers of holes that did not persist ((23)), and the number and distance threshold of those holes flexibly depended on attention and other cognitive processes. Our observations support the idea that there is information that can be found in features that do not persist ((24)).

We therefore summarize the topology of the correlation matrix as the peak Betti number, which is the maximal number of holes of a given type (called homology group) that appeared at any threshold (Figure 1D, (23)). We focus here on holes that are equivalent to circles (those detected by the first homology group) and spheres (detected by the second homology group), because these can be estimated using data sets of experimentally tractable size ((25); (26); (16)). For simplicity, we refer to these as circular and spherical features respectively. In our data and models, focusing on the peak Betti curve led to qualitatively similar conclusions as other common topological summaries (see similar conclusions in (27); (28); Supplementary Figures 1 and 2).

Here, we demonstrate that topological descriptions of correlated variability are an effective bridge between behavioral, physiological and theoretical approaches to studying neuronal populations. The peak Betti number is flexibly modulated by cognition, is related to performance on a visually guided task, and gives novel insights into mechanistic models and the function of real and artificial neural networks in different cognitive conditions.

### B. Topology as a bridge to behavior

The primary reason for focusing on noise correlations is that the magnitude of noise correlations in visual cortex has been strongly linked to performance on visually guided tasks ((5), (29)). To justify our use of persistent homology to study neuronal networks, we tested the hypothesis that topological signatures of network activity capture key properties of the relationship between correlated variability and behavior.

Four observations suggest that the peak Betti number captures the aspects of noise correlations that are related to performance. First, across recording sessions, there was a negative relationship between peak Betti number and the average noise correlation (Figure 2C, D)), meaning that sessions in which the average noise correlation was low tended to have a higher peak Betti number. Second, consistent with the observation that attention reduces noise correlations ((5); (30); (31); (2)), attention changes the peak Betti number (Figure 2A, B). Third, the peak Betti number was higher on trials in which the animal correctly detected a change in a visual stimulus compared to trial in which the animal missed the stimulus change (Attended condition average peak Betti Number *H*_1_ correct: 14.47,incorrect: 13.46;*H*_2_ correct:8.38,incorrect:7.41; Paired T test (peak Betti Number *H*_1_) *p* = 0.014, (peak Betti Number *H*_2_) *p* = 0.015). Finally, there was a positive correlation between peak Betti number and behavioral performance (Figures 2 E, F). Together, these results show that peak Betti number is a good description of the aspects of correlated variability that correspond to changes in behavior.

**Fig. 2.**
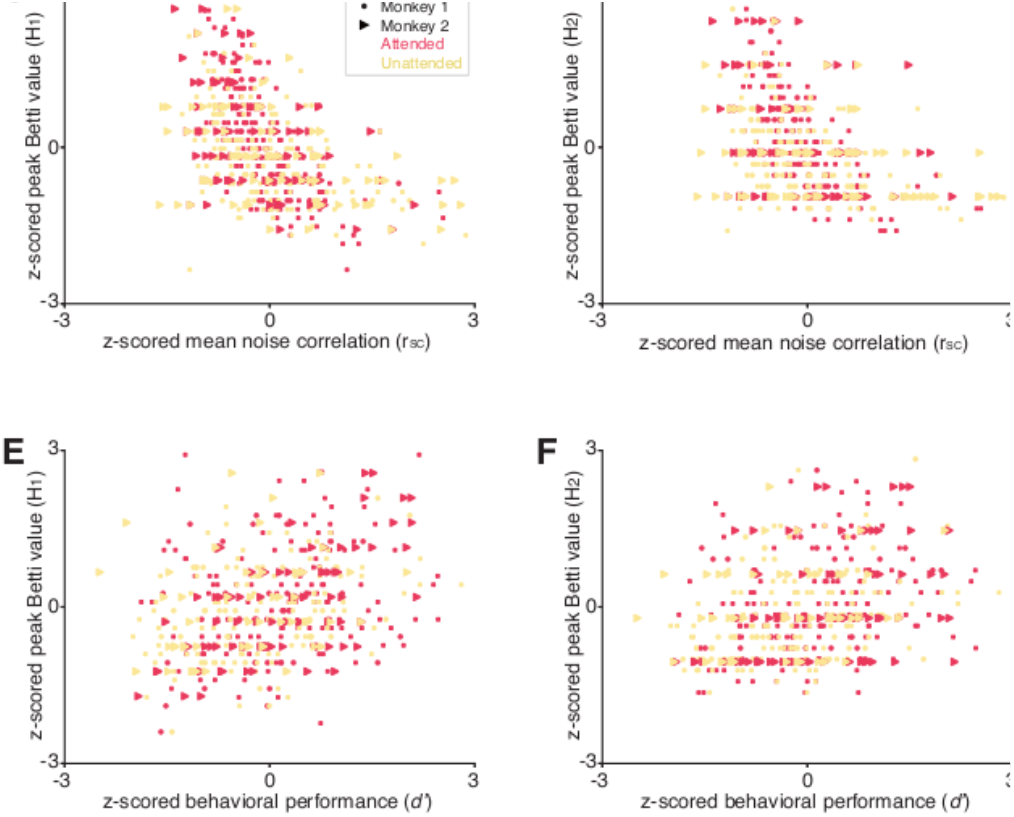
TDA reveals the relationship between neurons and behavior. (A, B) Attention increases the maximum number of circular features (i.e. 1*st* dimension features; A) and spherical features (i.e. 2*nd* dimension features; B) over the range of threshold values. Each point represents one experimental session, and the red points are the mean values, which are significantly greater for the attended condition (y-axes) than the unattended condition (x-axes; paired T-tests, p<0.01). (C) The maximum number of circular features is correlated with the mean *r_SC_* in both attention conditions both are z-scored for each animal; (Attended: *r* = −0.52, *p* = 1.3*e* – 18; Unattended: *r* = −0.42, *p* = 7.9*e* – 12; Paired T-test (Attended and Unattended, peak of the Betti curve):*p* = 8.41*e* – 5) (D) Same, for the maximum number of spherical features 2*nd* homology group (Attended: *r* = −0.44, *p* = 4.9*e* – 13; Unattended: *r* = −0.36, *p* = 5.3*e* – 9; Paired T-test (Attended and Unattended, peak of the Betti curve):*p* = 3.09*e* – 6) (E) There is a strong relationship between the maximum number of circular features and behavioral performance (*d*’ or sensitivity calculated for a single orientation change for each session; both measures are z-scored; Attended: *r* = 0.32, *p* = 2.34*e* – 7; Unattended: *r* = 0.3, *p* = 1.41*e* — 5) (F) Same, for the maximum number of spherical features (Attended: *r* = 0.27, *p* = 2.06*e* – 5; Unattended: *r* = −0.28, *p* = 4.67*e* – 5).

### C. Topology as a bridge to mechanism

The magnitude of correlated variability places strong constraints on circuit models of the neuronal mechanisms underlying attention (Figure 3A; (9); (10)). In particular, network models are constrained by the observation that attention changes correlated variability in essentially a single dimension of neuronal population space in area V4 (Figure 2, (32); (33); (10); (7)).

**Fig. 3.**
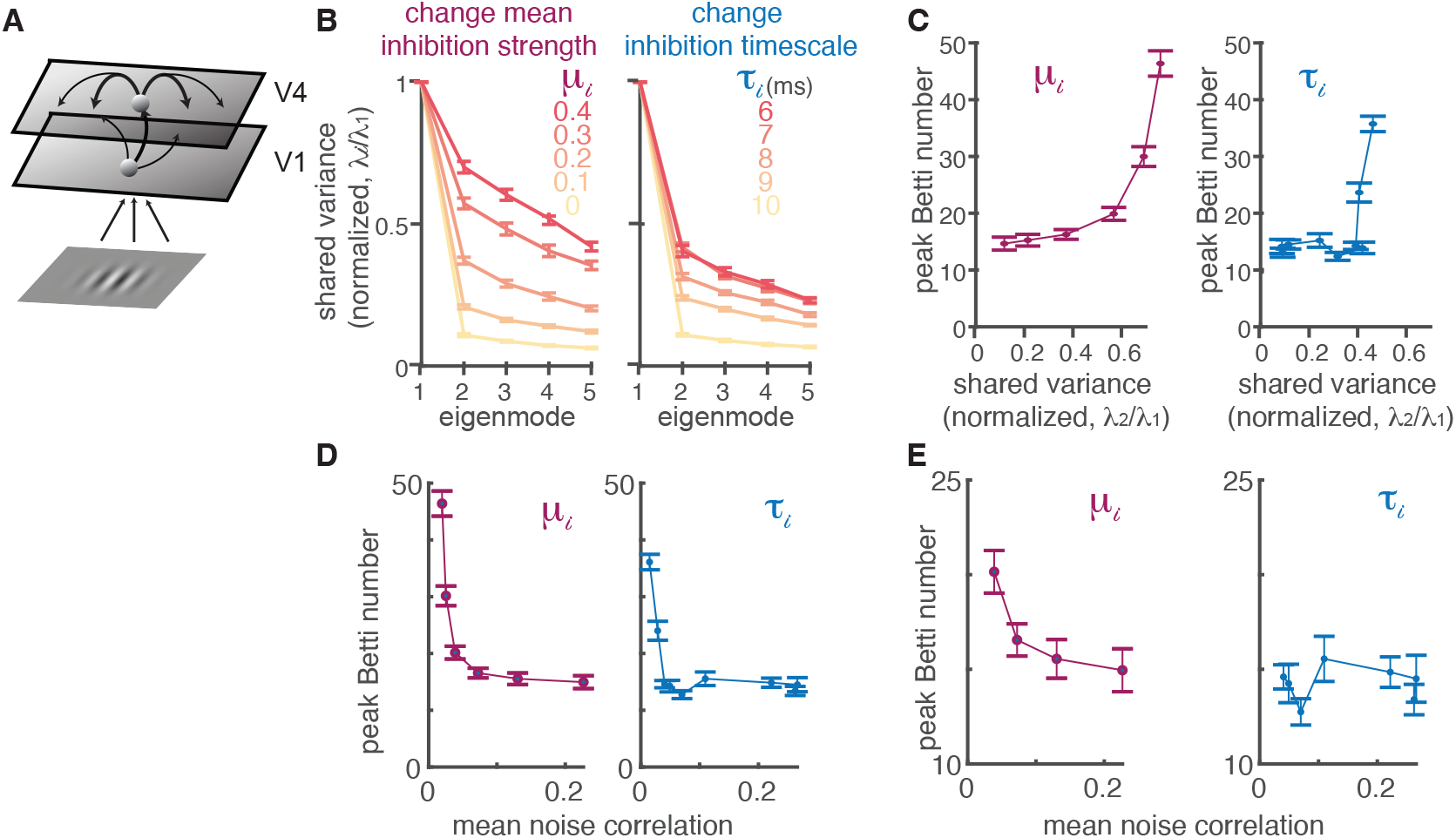
Figure 3: TDA can distinguish between different mechanistic models of attention. (A). Model schematic of a two-layer network of spatially ordered spiking neurons modeling primary visual cortex (V1) and area V4, respectively. The visual inputs to the model are the same Gabor stimuli used in our experiments. (B). Two distinct attention mechanisms can decrease correlated variability in a low rank way that is similar to linear descriptions of our data. We can reduce correlations either by increasing the currents to all inhibitory neurons (*μ_i_*) or decreasing the decay timescale of inhibitory currents 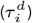. The plots depict the shared variance in each mode (the top five eigenvalues from the shared covariance, with private variance removed using Factor analysis), normalized by the shared variance in the first mode, for different values of *μ_i_* (left) or 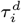 (right; error bars represent standard error of the mean; SEM). The two mechanisms appear indistinguishable using linear methods. (C). The two mechanisms cause different changes in the topological descriptions of the modeled V4 populations. As *μ_i_* increases (left panel), so does the shared variance present outside the first mode (x-axis) as well as the peak Betti number (shown for the circular (i.e. 1*st* dimension) features in the y-axis). Changes in *τ_i_* (right panel) result in a different relationship between the peak Betti number and the shared variance in higher dimensions, affecting the peak Betti number only at very short (unrealistic) timescales (those with the greatest shared variance; red lines in B). The peak Betti number is computed from the same simulated responses as in (B). Error bars represent SEM. (D) The peak Betti number (y-axis) has a different relationship with average noise correlation (x-axis) when modulated by changing the mean current to the inhibitory neurons (*μ_i_*, left panel) or decreasing the decay timescale (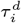, right panel). Error bars represent SEM. (E) Same as (D), except zoomed in to exclude parameter values that result in an unrealistically low mean noise correlation (<0.03). In this physiologically realistic range, changing *μ_i_* is associated with monotonic changes to the peak Betti number, while changing 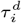 does not appreciably change peak Betti number.

Topological descriptions of simulated networks can distinguish between competing models in situations when the magnitude of shared variability, even in the most relevant dimension, fails to do so. We analyzed the outputs of our spatially extended network of spiking neuron models, which internally generate correlated variability through spatiotemporal dynamics ((10)). In the model, the magnitude of correlated variability can be changed by modulating inhibition in two distinct ways: either increasing the input drive to the inhibitory neurons (*μ_i_* in Figure 3B) or decreasing the timescale of inhibition (*τ_i_* in Figure 3B) changes correlated variability in a low rank way.

These two mechanisms have very different effects on the topology of the correlated variability, even when the mean variability is equivalent. For most parameter values, changing the input drive to the inhibitory neurons has a much greater effect on the peak Betti number than changing the timescale (Figure 3C, D, E). While changing the timescale of inhibition is extremely common in circuit models (for review, see (10)). in real neural networks, the timescale of inhibition is longer than excitation and is inflexible ((34); (35); (36)). Both the biology and the topology are consistent with the idea that attention instead acts by increasing the input drive to the inhibitory neurons ((9); (10)).

These results demonstrate that topological signatures of correlated variability provide constraints on mechanistic models that are unavailable using linear measures of neural activity. Changes in the mean or dimensionality of correlated variability are not necessarily coupled with changes in the topological signatures of the network. Together, our results highlight the value of using circuit models as a platform on which to test and generate hypothesized mechanisms underlying perception and cognition.

### D. Topology as a bridge to network function

The past two decades have seen an explosion in the number of studies demon-strating that correlated variability depends on a wide range of sensory, cognitive, and motor conditions that change behavior (for review, see (2)). Despite much effort from the experimental and theoretical neuroscience communities ((33); (37); (10); (9); (32); (38); (39)), how changes in correlated variability might affect behavior remains unclear. The observations that topological summaries of the noise correlation matrix are related to behavior suggest that, via known connections to network control theory ((40); (41); (42)), TDA can provide insight into the relationship between correlated variability, the function of a network, and behavior.

Topological data analysis has known connections to network control theory because both measure the structure (or lack thereof) in a functional connectivity matrix ((42)). We reasoned that network control theory, which seeks to quantify the ability of interventions (in our case, visual stimuli, cognitive processes, or random fluctuations) to alter the state of a network ((43)) could provide intuition about the relationship between TDA and the function of our neuronal network. While network control theory is primarily used in engineering, recent work has used controllability to quantify the flexibility of large neural systems, constrained by fMRI data ((44)).

These methods focus on quantifying the energy required to move between states of the neural population. We define a state as the vector of neural population activity on a given trial, and the energy required to move between states is constrained by the noise correlation matrix (e.g. in Figure 4A). For example, if the responses of all the neurons are highly positively correlated, then reaching a state in which the response of some is high while the others are low is unlikely and therefore requires significant energy.

**Fig. 4.**
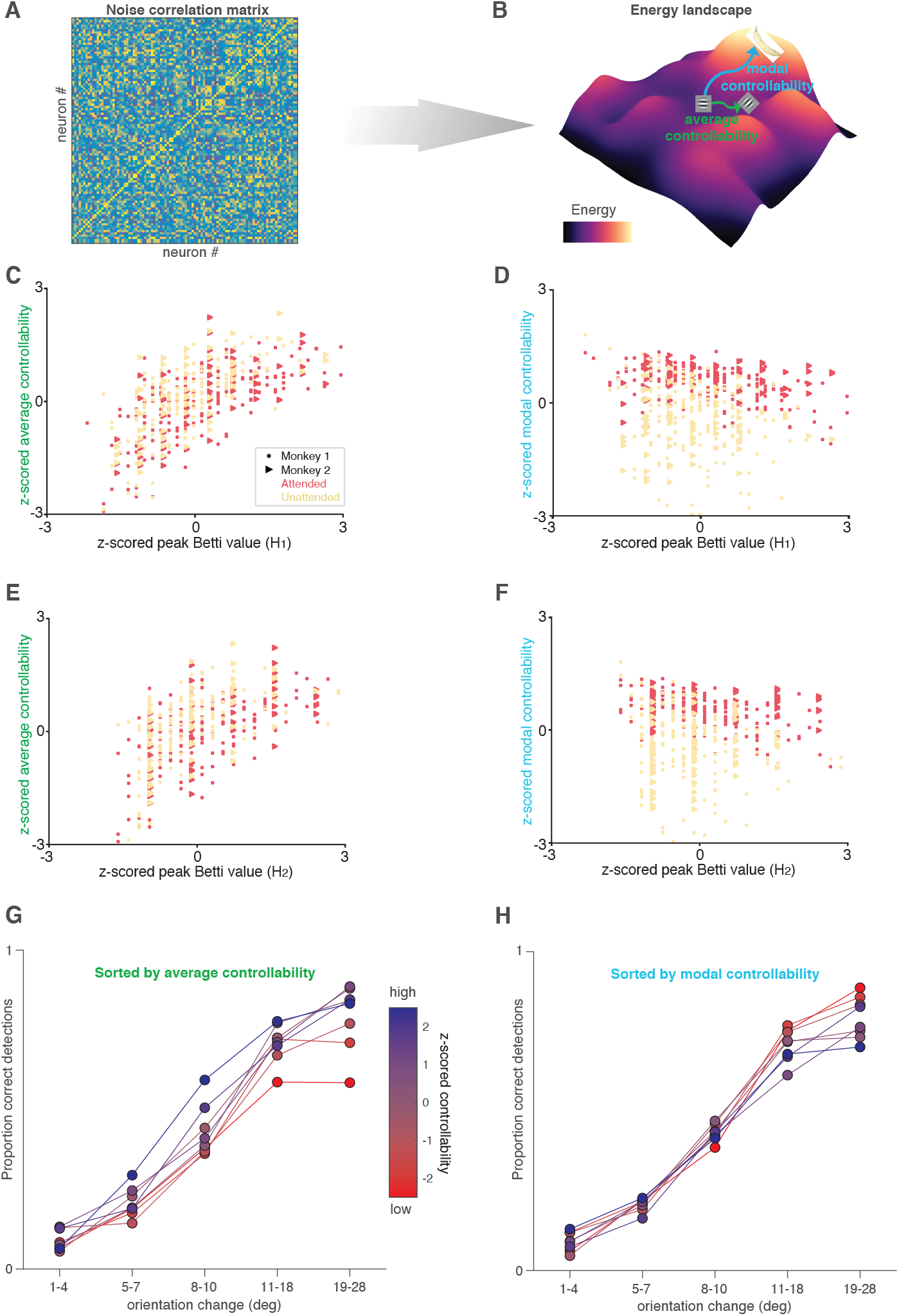
TDA and controllability provide new insight into network function. (A,B). Illustration of our controllability calculation. We consider the noise correlation matrix (A) as a functional connectivity matrix, and use this to calculate an energy landscape (illustrated for a hypothetical situation in B; colors indicate energy). Average controllability is defined as the energy required to move from a starting point (e.g. a response to a horizontal Gabor stimulus) to nearby states (e.g. a response to an oblique Gabor), and modal controllability is defined as the energy required to move to distant states (e.g. thinking about a banana). (C). High average controllability is associated with maximum number of circular (i.e. 1st dimension) features (Both measures were z-scored for each animal, and the lines were fit for each attention condition; attended: *r* = 0.65, *p* = 1.03*e* – 31; unattended: *r* = 0.68, *p* = 1.28*e* – 33; Paired T-test (Attended and Unattended, average controllability):*p* = 7.5*e* – 61). (D) High modal controllability is associated with lower number of circular (i.e. 1st dimension) features (Conventions as in A; (attended: *r* = −0.37, *p* = 7.7*e* – 10; unattended: *r* = –0.23, *p* = 3.2*e* – 4; Paired T-test (Attended and Unattended, modal controllability):*p* = 1.5*e* – 38) E) Relationship between average controllability and maximum number of spherical features (i.e. 2nd dimension) (attended: *r* = 0.59, *p* = 3.5*e* – 25; unattended: *r* = 0.59, *p* = 9.5*e* – 24). Conventions as in (C). (F). Relationship between modal controllability and number of spherical (2nd dimension) features (attended: *r* = −0.38, *p* = 3.12*e* – 10; unattended: *r* = −0.18, *p* = 4.7*e* – 3). Conventions as in (D). (G). High average controllability is associated with better performance at all orientation change amounts. Colors represent z-scored average controllability (the experimental sessions were split into six equally sized bins by average controllability) and the plot shows proportion correct detections (hit rate) as a function of orientation change amount. (H). High modal controllability (bluer colors) is associated with a worse lapse rate (worse performance on easier trials). Conventions as in (H).

If our starting point (e.g. the horizontal Gabor in Fig. 4B) is the population response to a horizontal Gabor stimulus presented before the orientation change in our task (see Fig. 1A), a nearby state might be the population response to the changed stimulus (e.g. the oblique Gabor in Fig. 4B). A distant state might be a population response when the monkey is concentrating on something very different and task-irrelevant (e.g. thinking about the banana in Figure 4B). Average controllability quantifies how readily the population moves from the starting point to nearby states while modal controllability quantifies how readily the population moves to distant states.

There is no mathematical relationship between average and modal controllability. Indeed, average and modal controllability were uncorrelated across sessions in our data (R=0.008; p=0.9).

However, we discovered that the topological descriptions of neuronal population are strongly related to both average and modal controllability, and both are related to attention. High peak Betti value (which occurs more readily in the attended state) is associated with decreases in the energy required to drive the system to nearby states (high average controllability; Fig. 4C, E). In contrast, there is a negative relationship between peak Betti value and the energy required to drive the system to distant states (modal controllability; Fig. 4D, F) but changing attention conditions increases modal controllability (compare the red and yellow points in Fig. 4D, F).

The different relationships between topology and average and modal controllability observations give new insight into the tradeoffs associated with attention. Attending to a stimulus improves the network’s ability to respond to subtle interventions, which is consistent with the attention-related improvements in the animal’s ability to detect a subtle change in the visual stimulus (Fig. 1A), but it has complex effects on the ability of the network to change states dramatically, which may mean that attention reduces cognitive flexibility. In future work, it would be interesting to study whether changes in controllability can account for change blindness and other behavioral demonstrations that attention reduces the ability of observers to notice very unexpected stimuli, such as the classic example of failing to notice a gorilla walking through a basketball game ((45)).

Indeed, average and modal controllability have distinct relationships with the monkeys’ performance in our task. We sorted the experimental sessions by average controllability (colors in Fig. 4G) or modal controllability (colors in Fig. 4H). Increased average controllability was associated with improvements in the monkeys’ ability to detect all orientation change amounts (except the smallest changes in which a floor effect meant that they were rarely detected). In contrast, modal controllability was unrelated to the monkeys’ ability to detect subtle orientation changes and was anticorrelated with the ability to detect large, easy orientation changes. One interpretation is that when modal controllability is high, the monkeys’ minds wander more easily to distant, potentially task-irrelevant states, increasing the lapse rate on easy trials. Together, these results demonstrate that in addition to linking to behavior and mechanism, topological signatures of the structure of noise correlations can provide insight into the function of the network and the behavioral trade-offs associated with changes in correlated variability.

### E. Comparing topological summaries and mean noise correlations

The results in Figures 1–4 demonstrate that topological summaries of noise correlation matrices are related to many quantities of interest, including behavior, average and modal controllability, and the mean pairwise noise correlation. If all of these quantities are related to each other, what is the added value of TDA over the simpler and more common mean noise correlation metric?

Others have written about the value of TDA for many applications, including analyzing signals from groups of neurons (as opposed to noise as we have done here; (22)). Ours results suggest two key advantages of TDA over mean noise correlations for understanding the mechanisms by which populations of neurons guide behavior. First, we demonstrated that TDA can distinguish between models in which the mean noise correlations (and even the dimensionality of the noise correlation matrix) were indistinguishable (Fig. 3).

Second, we found that the relationships between topological summaries and other quantities of interest (including performance on our change detection task, average controllability, and modal controllability) are stronger and dissociable from the relationships between those quantities and mean noise correlations (Fig. 5). To established this, we first computed the raw session-to-session correlation coefficient between each quantity of interest (Fig. 5A, D, which are identical except that Fig. 5A contains the peak Betti value in the first dimension as a representative topological summary and Fig. 5D contains mean noise correlations). Next, we computed partial correlations between those same quantities while controlling for the effect of mean noise correlation (Fig. 5B) or the peak Betti value (Fig. 5E). The difference between the raw and partial correlations reflects the extent to which, for example, the relationship between behavior and average controllability, can be attributed to the relationship between each of those and mean noise correlation (Fig. 5C) or the peak Betti value (Fig. 5F).

**Fig. 5.**
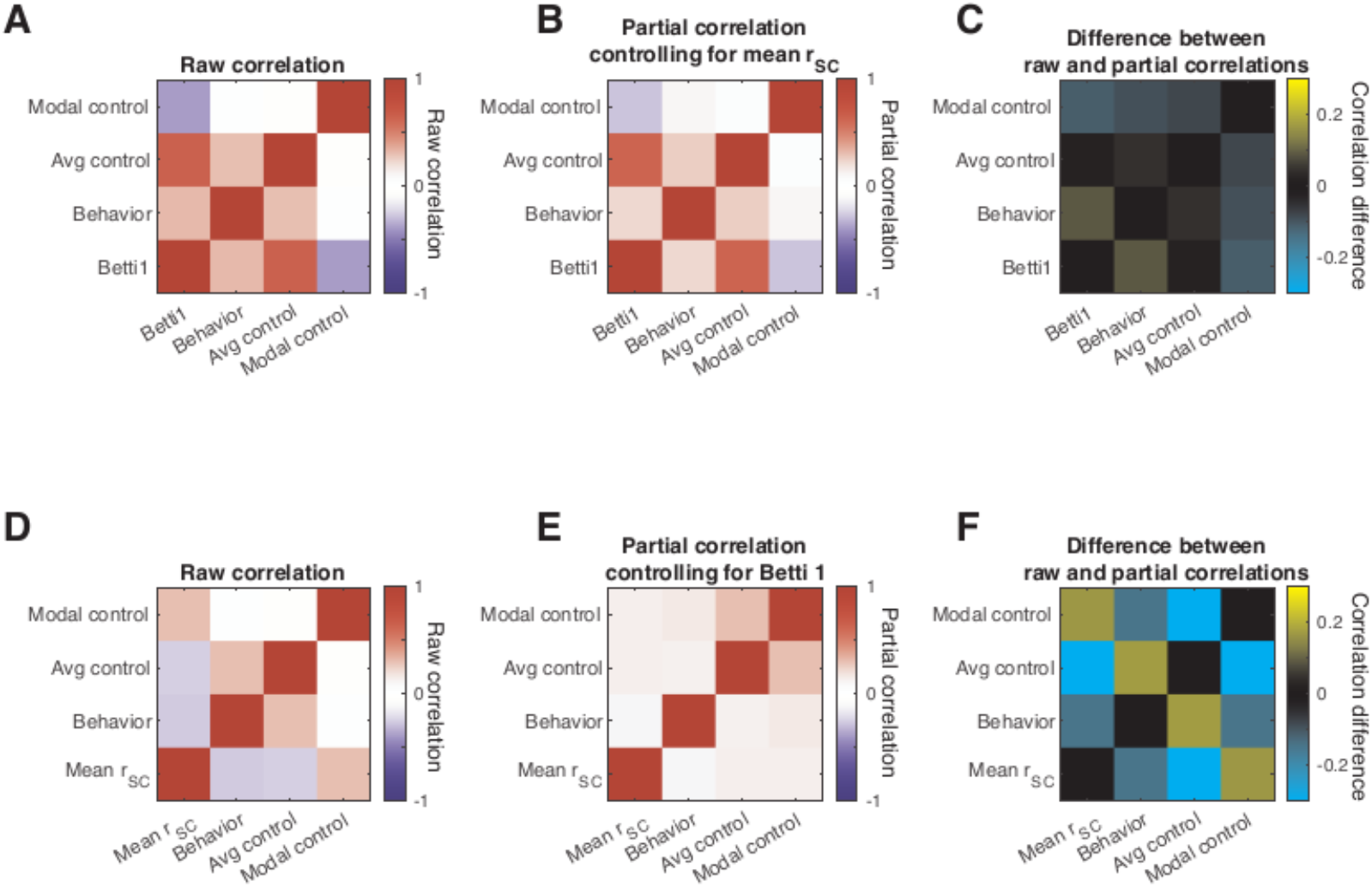
A partial correlation analysis reveals that topological summaries of noise correlation matrices provide insights that are distinct from the mean noise correlation. A. Raw correlation coefficients summarizing the session-to-session correspondence between modal controllability, average controllability, behavior (defined as perceptual sensitivity as in Fig. 2), and the peak Betti value (maximum number of circular features). The relationships between the controllability and behavioral metrics and other topological descriptions (including the maximum number of spherical features and total persistent) are qualitatively similar, so for simplicity, we focus on the maximum number of circular features here. Diagonals represent self-correlations, and are therefore 1 by definition. B. Same as A, but the colors reflect partial correlations that control for session-to session-variability in mean noise correlation. C. Difference between A and B, showing that the raw and partial correlations are qualitatively similar. D. Same as A, but the last row is mean noise correlation instead of the peak Betti value. E. Partial correlations controlling for session-to-session variability in peak Betti value. F. Difference between D and E.

Controlling for the peak Betti value had bigger impact on every pairwise relationship than controlling for mean noise correlation (and this difference was statistically significant for all three common off diagonals in Fig. 5C and 5F; p<0.05 with a Bonferroni). This result indicates that statistically speaking, the peak Betti value provides more and independent insight into behavioral and control theory measures of circuit function than mean noise correlation.

It is worth noting that in all likelihood, none of the measures discussed here (e.g. from TDA, control theory, or other descriptions of the noise correlation matrix) are quantities that are directly used by the brain in neural computations. Neural computations are performed to guide behavior at individual moments or on individual trials, and noise correlations, or any derivative of them, are computed over many ostensibly identical trials. The value of any of these metrics is that they provide insight into the underlying computations. Our results (especially in Fig. 5) demonstrate that TDA provides insight into some key quantities (perhaps most importantly into behavior) that are distinct from the insights that can be gleaned from mean noise correlations alone.

## 2. Discussion

### A. Implications for topology

Although TDA has been used for many scientific applications ( (18)), our use of TDA differs substantially from previous work. The prevailing paradigm used in virtually all TDA applications, including many in neuroscience ( (17); (13)), focuses on identifying persistent topological features, such as holes that persist across many thresholds (Figure 1B). These persistent features are appealing because they can reveal the structure of a simple network. However, applying these methods to analyze neural circuits may not lead to any scientific discoveries, since persistent features are not expected in neural response variability, which is thought to arise from complex network properties that make it relatively unstructured ((4)).

However, we demonstrated that using TDA to analyze correlated variability in neuronal responses is useful, even in the absence of persistent features. The link that we demonstrated between the topology of noise correlations, which have been shown to reflect both cognition and the anatomy of the system ((3)), and the controllability of the network on individual trials (which are what matter for guiding behaviors) therefore has implications far beyond neuroscience. Throughout the natural and physical sciences, natural systems are complex and call for sophisticated data analytics pipelines. In astronomy, for example, TDA has been used to understand the relationship between planets, stars, and galaxies on a huge range of spatial scales. Our use of TDA to analyze non-persistent topological features (see also (23)) will be a bridge between neuroscience and other fields. These tools for analyzing and interpreting complex networks can be deployed in many other scientific domains.

### B. Implications for neuroscience

We demonstrated here that using TDA to analyze the variability neural populations can illuminate novel links between behavior, neurons, computations, and mechanisms. This sort of bridge between different levels of investigation has the potential to be broadly transformative. In an age of massive improvements in experimental technologies and tools for measuring the activity of large numbers of neurons, perhaps the greatest barrier to success understanding the neural basis of behavior is that it is different to compare and integrate results from experiments using different methods in different model systems. TDA can reveal relationships between neural networks, computations, and behaviors that are robust to the differences in neuronal responses that occur between every different experimental system ((46)). These analytical links make it possible to leverage the complementary strengths of each approach.

A holistic view of neuronal populations is necessary for understanding any neural computation. Essentially every normal behavioral process or disorder of the nervous system is thought to involve the coordinated activity of large groups of neurons spanning many brain areas. Tools for understanding and interpreting large populations have lagged far behind tools for measuring their activity. Standard linear methods have provided a limited view, and the field is in dire need of a new, holistic window into population activity. Our results demonstrate a hopeful future for using the topology of neural networks to fulfill that need.

## Materials and Methods

### C. Experimental methods

Different analyses of these data have been presented previously ((5)). Briefly, two adult rhesus monkeys (Macaca mulatta) performed an orientation change detection task with a spatial attention component ((30)). The monkeys fixated a central spot while two peripheral Gabor stimuli flashed on (for 200 ms) and off (for a randomized period between 200 and 400 ms). At a random and unsignaled time, the orientation of one stimulus changed, and the monkey received a liquid record for making an eye movement to the changed stimulus within 500 ms. We cued attention in blocks of 125 trials, and the orientation change occurred at the cued location on 80% of trials. Our analyses are based on responses to the stimulus presentation before the change, which was the same on every trial within a recording session. The location, contrast, and spatial frequency of the Gabor stimuli were the same during every recording session, but the orientation differed across sessions. The location of one stimulus was within the receptive fields of the recorded neurons and the other stimulus was in the opposite hemifield.

We presented the stimuli on a CRT monitor (calibrated to linearize intensity; 120 Hz refreshed rate) placed 52 cm from the monkey. We monitored the animals’ eye position using an infrared eye tracker (Eyelink 1000; SR Research) and recorded eye position, neuronal responses (30,000 samples/s) using Ripple Hardware.

While the monkey performed the task, we recorded simultaneously from a chronically implanted 96-channel microelectrode array (Blackrock Microsystems) in the left hemisphere of visual area V4. We include both single units and sorted multiunit clusters (mean 34 and 15 units for Monkeys 1 and 2, respectively). The average number of simultaneously recorded pairs of units (for computing noise correlations) was 561 for Monkey 1 and 105 for Monkey 2. The data presented are from 42 recording sessions from Monkey 1 and 28 recording sessions from Monkey 2.

### D. Data preparation

To examine how the topology of networks of neurons in visual cortex or outputs of spiking models depend on attention, we constructed distance matrices from noise correlation matrices ((3)). We defined the noise correlation for each pair of neurons (also known as spike count correlation; (3)) as the correlation coefficient between the spike count responses of the two neurons in response to repeated presentations of the same stimulus. We based our analyses on spike count responses between 60-260 ms after the onset of the visual stimulus to allow for the latency of visual responses in area V4. We used responses to the stimulus before the orientation change because those are the same on every trial. We focused on trials when the monkey correctly identified the changed stimulus and compared responses in the two attention conditions.

Many measures of neuronal activity depend on experimental details like the number of recorded neurons or their mean firing rates, which were different for the two monkeys (see (5) for details). To allow us to combine across animals, we z-scored the results for each animal (across both attention conditions) and plot those normalized measures in the figures.

### E. Behavioral measures

To analyze the relationship between neuronal responses and behavior, we adopted a signal detection framework ((47); (48)) to assess how behavior depends on neurons and attention ((49);(50); (51); (52); (53); (54); (55)). Criterion is defined as

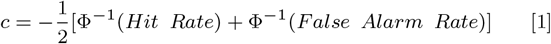

where Φ^-1^ is the inverse normal cumulative distribution function. Negative values of c indicate that the subject has a liberal criterion (bias toward reporting changes), and positive values indicate a conservative criterion (bias toward reporting nonchanges).

Sensitivity is defined as

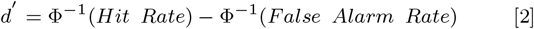

Larger values of d’ indicate better perceptual sensitivity.

Different orientation change amounts were used in different recording sessions. To compare across sessions, we fit the psychometric curve using a Weibull function and computed performance at a single, fixed orientation change for each session (see Figure 1A and (5)).

### F. Topological measures

We examine the topology of the noise correlations in each attention condition using a Vietoris Rips construction. This consists of defining a distance matrix (better understood as a weighted adjacency matrix), which we constructed from the noise correlation matrix *r_SC_*, to define pairwise (and higher order) connections between the vertices (representing neurons) in the simplicial complex. The distance matrix was chosen to be 1 – *r_SC_*, so that higher weighted interactions (i.e. those neurons which are strongly correlated) are defined as shorter distance and therefore entered the simplicial complex first. For brevity, we refer to this matrix as the distance matrix of the space while acknowledging that our measure does not satisfy the axioms of a distance function.

We consider a distance threshold which defines those pairwise interactions that are permitted to be considered in the simplicial complex. Such a process allows us to examine the evolution of the simplicial complex across different distance thresholds. We assess several properties of the simplicial complex at each threshold value, including the existence of holes (or higher dimensional voids) of a given dimensionality (termed homology dimension). A hole signifies a lack of connections (i.e. differences in the degree of correlation) between a subset of neurons at the current distance threshold. We focus our analysis on the first and second homology dimensions (which correspond to holes that are topologically equivalent to circles and spheres, respectively) because they can be estimated reliably given the size of our data sets.

In most applications of TDA, researchers focus on persistent features that imply a nontrivial structure. For example, imagine a set of people seated around an oval shaped table. We could use as a distance metric the physical distance between them, and we would consider two people to be ’connected’ if they are sitting closer than some distance threshold. At very small distance thresholds, no pair of people would be close enough to be considered connected. At very large thresholds (e.g. longer than the length of the room), all pairs of people would be connected. But for a large range of intermediate distance thresholds, each person would be connected to at least their nearest neighbor but would not be connected to everyone else, and the resulting graph would have a ’hole’, corresponding to the center of the table. In TDA, this hole would represent a persistent feature and would indicate that the seating arrangement has a particular structure. The presence of a single, persistent hole in the first homology dimension (equivalent to a circle), would imply that the people had arranged themselves around a table, but it would not specify whether that table was a circle, a rectangle, or another topologically equivalent shape.

Next, consider a situation where the same people simply sat in a haphazard arrangement on the floor. By chance, there would at some distance thresholds be holes around which, for example, a subgroup of people were arranged. Those chance holes would not persist for a very long range of distance thresholds. An intermediate situation, in which there is some structure to the seating arrangement (e.g. people sitting in small clusters), might have a smaller number of holes that persist for only a small range of thresholds.

Some recent studies have used TDA to investigate the signals encoded by populations of neurons or brain areas ((19);(20); (21); gardner2022toroidal). In these studies, the vertices represent trials, stimuli, or time periods, and the authors construct a distance metric to relate population responses at those different times. Because neural signals have structure, those authors were able to analyze features that persist over a long range of distance thresholds.

Our approach was orthogonal. We took each neuron to be a vertex, and the distance between them was given by the pairwise noise correlation. Noise correlation matrices are in general thought to be unstructured, which is why most previous studies focus on their mean or linear dimensionality ((3), (10)). Consistent with this idea, we did not observe notable persistent features (see Figure 1C for an example of observed features). Therefore, the ’holes’ in our data should be thought of as topological noise. The relationship between this topological noise and other quantities of interest (e.g. behavior) indicates that although no individual hole is particularly important, the distribution of them can provide insight into neurobiological processes.

We therefore adopted the approach of focusing on the distribution of topological features rather than on looking for long persistent cycles ((24)). We examined how properties of the generated Betti curves (such as the peak or total persistence) relate to common measures of attention like average noise correlations and behavioral performance.

### G. Topological data analysis example

We provide here a detailed walk-through of the schematic in Figure 1B. Suppose that we have a group of vertices. You can think of these vertices as a collection of neurons where each vertex is a neuron. Indeed, this is the view that we take in this work. Along with this group of vertices we have an underlying weight matrix that expresses not only what vertices are connected to one another but the strength of these connections. A neuroscience interpretation of this weight matrix is the connectivity matrix of a population of neurons. If we look at a pair of vertices, say *n*_1_ and *n*_2_ and observe an entry of 0.5 in the weight matrix, then we know that not only are *n*_1_ and *n*_2_ connected but the strength of their connection is 0.5.

With both a group of vertices and a weight matrix expressing how the vertices are connected, we can now apply topological methods. These methods will allow us to examine relationships between subgroups of vertices. We define a threshold value that will determine which connections in the weight matrix are allowed. Allowed connections are those whose value is at most the threshold value. However, given that the “optimal” threshold value is unknown, the typical approach is to vary the threshold value over a range while simultaneously tracking the properties of the evolving graph. We choose to track the number of holes (i.e. a empty space in the graph due to a lack of connections). As we continuously increase the threshold, certain connections will enter the graph and the connections that are lacking may form holes within the graph.

We examine this process in the context provided in Figure 1B. Initially there are 7 disconnected vertices. We can assume that this is the case because the distance threshold is 0 at the beginning. As we increase the distance threshold, we have 1 isolated vertex and a ring structure. Given that there is a lack of connections between different subsets of these vertices, a hole is formed. Thus, the number of holes for our structure (i.e. referred to as the Betti Number in the literature) is 1. As we increase the distance threshold further, the isolated vertex becomes connected to 2 vertices and we signify this all-to-all connected sub-graph by a shaded region. Observe that although the distance threshold has increased, the hole in the center of the ring is still present and thus our Betti Number is still 1. Finally, we increase the threshold, and all vertices in the ring become connected. Our Betti number is now 0. This has occurred because the weights of those connections are at most the value of the distance threshold. We apply this same process to our neural data where the weight matrix is one minus the noise correlation value between a pair of neurons in the overall noise correlation matrix.

### H. Spatial model construction

Our spatial model is a variation of the two layer network of neurons discussed in ((10)). Neurons in this network are arranged uniformly on a [0, 1]×[0, 1] grid. The first layer (i.e. the feed-forward layer) consists of *N_x_* = 2500 excitatory neurons that behave as independent Poisson processes. The second layer consists of 40, 000 excitatory and 10, 000 inhibitory neurons that are recurrently coupled. The second layer receives input from the first layer. The network’s connectivity is probabilistic but dependent on a Gaussian of width *σ*\*. Thus neurons that are further away from each other on the grid are less likely to connect.

The parameters are the same as the two-layer network in the Huang et al., 2019 and are chosen to approximate known biology of cortical circuits (see (10) for details). Specifically, the synaptic strengths are scaled by 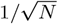, where *N* is the total number of neurons in the network, as used in the so-called balanced networks ((56)) such that the recurrent network can internally generate variability in neural spiking for large *N*. The projection widths of the excitatory and inhibitory neurons are chosen to be the same (*σ_e_* = *σ_i_* = 0.1), which is consistent with anatomical findings from visual cortex ((57); (58)). Each neuron is modeled as an exponential intergrate-and-fire neuron model, following standard formulation in past work ((59)).

The only differences between the published model and one here are the following. The feedforward connection strength from layer 1 to layer 2 is *J_ex_* = 140 and *J_ix_* = 0 for excitatory and inhibitory neurons, respectively. Fig 3B left: *μ_i_* = 0, 0.1, 0.2, 0.3, 0.4 and *τ_i_* is 10. Fig 3B right: *μ_i_* = 0, and *τ_i_* = 6, 7, 8, 9, 10 ms. These parameters were chosen so that the manipulations of *μ_i_* and *τ_i_* begin with the same parameter set at high correlation value, which is unstable with turbulent wave dynamics. There were a total of 15 simulations of 20 seconds each for each parameter conditions. The first 1 second in each simulation was removed. The spike counts were computed using 140 ms time window to mimic the data. x

We implemented this model using EIF neurons. The voltage dynamics of these neurons are governed by the following equation((10)):

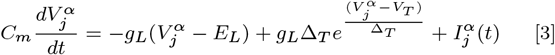

where 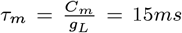, *E_L_* = −60*mV*, *V_T_* = −50*mV*, *V_th_* = −10*mV*, Δ_*T*_ = 2*mV, V_re_* = −65*mV*, *τ_ref_* = 1.5*ms* and the total current 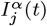 obeys the following equation:

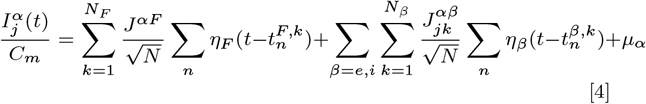

where *N* is the total number of neurons in the second layer and *μ_α_* is the static current to the *α*(∈ {*E, I*}) population. *η_β_* is the postsynaptic current given by the following equation

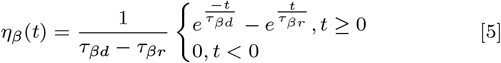

where the rise time constant *τ_βr_* = 5. We consider multiple values of the decay time constant *τ_βd_*. For both dimensionality (see Factor Analysis section) and topological (see Topological Measures section) comparisons, we also considered a range of values of the *μ_I_* parameter which correspond to the overall depolarization of the inhibitory population and which has been shown to affect the dimensionality of the generated data.

### I. Factor analysis

To assess the dimensionality of the population simulated using the spatial model with different parameters, we used factor analysis ((39)). We based our analysis on a number of neurons by number of trials matrix of spike counts of the simulated excitatory neurons. We then used that matrix to compute a spike count covariance matrix. Factor analysis separates the spike count covariance matrix into a shared component that represents how neurons co-vary together and an independent component that captures neuron-specific variance. Following the notation of ((39)), use *L* to refer to the loading matrix relating m latent variables to the matrix of neural activity. In this way, the rank, m, of the shared component *LL^T^* is the number of latent variables that describes the covariance. We refer to the independent component as Ψ, which is a diagonal matrix of independent variances for each neuron. We then assess the dimensionality of the network activity by analyzing the eigenvalues of *LL^T^*. To focus on dimensionality rather than the total amount of independent or shared variance (which depends on many model parameters), we normalized each eigenvalue by dividing by the largest eigenvalue.

We performed this analysis on the spike count responses of a randomly sampled 500 simulated neurons. All analyses were cross-validated. Error bars in the figures come from analyzing many instances of the network generated using fixed model parameters.

### J. Controllability measures

The goal of our controllability measures is to understand how the noise correlation matrix in each attention state constrains estimates of the function of the network. We consider a hypothetical (possibly nonlinear) dynamical system whose dynamics can be linearized and whose effective connectivity is defined by the noise correlation matrix. We analyze the properties of the system to assess the amount of effort it takes to change the system’s state using external input. We summarize these calculations using two standard measures of controllability ((60); (43)): average controllability, which relates to the ability to push the system into nearby states or states with little energy and modal controllability, which relates to the ability to push the system into distant states or states that require more energy.

We take the effective connectivity matrix A to be the noise correlation matrix generated from the spike count responses to repeated presentations of the same visual stimulus as described above. To align with the controllability methods in previous cognitive neuroscience studies (e.g. (61), (62); (60); (63)) and to remove the influence of self connections (which are defined as 1 for a correlation matrix), we set the diagonals of the effective connectivity matrix A to 0. (Leaving the diagonals as 1 did not qualitatively change our results).

We then consider the energy required to steer the network from an initial state *x*_0_ to a target state *x*(*T*) = *x_T_*.

The average controllability *ϱ_c_* is defined as

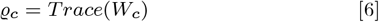

where 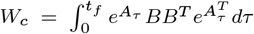 is the controllability gramian matrix in which *B* represents a matrix of nodes (neurons) in which we could inject hypothetical inputs to change the network state (the full matrix in our case) and *τ* represents the fact that the input could be in principle time varying. *Trace*() is the trace of the matrix (i.e. the sum of the diagonal elements of the matrix).

The modal controllability *ϕ_i_* is defined as

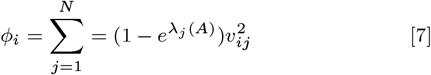

where *λ_j_* is the jth eigenvalue of the effective connectivity matrix *A*. *v_•j_* corresponds to the eigenvectors of the effective connectivity matrix *A*.

We used the equations given above to compute the average and modal controllability of the recorded population of neurons. To determine the relationship between controllability and behavior (Figures 4G,H), we computed average and modal controllability for each session, z-scored those measures for each monkey, and divided the sessions into six equally sized bins for each controllability measure.

## ACKNOWLEDGMENTS

We thank Douglas Ruff, Joshua Alberts, and Jen Symmonds for contributions to the previously published experiments and Alessandro Rinaldo and Ralph Cohen for helpful conversations. This work was supported by NIH grants 2R01EY022930 (MRC, TCR), RF1NS121913 (MRC, CH), R01EY034723 (MRC), and K99NS118117 (AMN), and grant 542961SPI from the Simons Foundation (MRC).

